# Limits of subjective and objective vection for ultra-high frame rate visual displays

**DOI:** 10.1101/2020.03.19.998591

**Authors:** Séamas Weech, Sophie Kenny, Claudia Martin Calderon, Michael Barnett-Cowan

## Abstract

Large-field optic flow generates the illusory percept of self-motion, termed ‘vection’. Smoother visual motion displays generate a more compelling subjective sense of vection and objective postural responses, as well as a greater sense of immersiveness for the user observing the visual display. Research suggests that the function linking frame rate and vection asymptotes at 60 frames per second (FPS), but previous studies have used only moderate frame rates that do not approach the limits of human perception. Here, we measure vection using subjective and objective (mean frequency and path length of postural centre-of-pressure (COP) excursions) responses following the presentation of high-contrast optic flow stimuli at slow and fast speeds and low and ultra-high frame rates. We achieve this using a novel rendering method implemented with a projector capable of sub-millisecond temporal resolution in order to simulate refresh rates ranging from very low (15 FPS) to ultra-high frame rates (480 FPS). The results suggest that subjective vection was experienced most strongly at 60 FPS. Below and above 60 FPS, subjective vection is generally weaker, shorter, and starts later, although this pattern varied slightly according to the speed of stimuli. For objective measures, while the frequency of postural sway was unaffected by frame rate, COP path length was greatest for 480 FPS stimuli. Together, our results support diminishing returns for vection above 60 FPS and provide insight into the use of high frame rate for enhancing the user experience in visual displays.

## Introduction

Observing large-field optic flow is sufficient to evoke the illusory perception that the body is in motion. This illusion, known as vection, provides a convincing example of the visual control of posture and gait (Delorme & Martin, 1986; Lishman & Lee, 1973). Vection is accompanied by postural reflexes that indicate the engagement of motor strategies that aim to maintain stable control of the body during perceived self-motion (Berthoz et al., 1979). Recent evidence suggests that postural excursions during vection hold strong predictive power in models that classify susceptibility to motion sickness at the individual level (Keshavarz et al., 2015; Weech et al., 2018).Several other insights have been provided by studies on vection, including the neural basis of self-motion perception (e.g., Brandt et al., 1998; Kovács et al., 2008; Wada et al., 2016) and the relationship between visual eccentricity and self-motion (Johansson; Post, 1988; Nakamura & Shimojo, 1998). Studies of vection have an applied focus: Recently, the possibility of improving virtual reality experiences using vection has been investigated by several groups (Nakamura et al., 2016; Riecke, 2010; Riecke et al., 2015; Weech & Troje, 2017). These studies suggest that generating a convincing illusion of self-motion is related to compelling visual experiences that enhance the ‘presence’ of the viewer (Prothero, 1998; Ohmi, 1998; Weech et al., 2019).

Advances in visual display technologies and recording methods have brought about an interest in the effect of high frame rate displays on visual experiences (Watson, 2013; Wilcox et al., 2015). While high frame rate video holds appeal due to the prospect of increased immersiveness, there are widely documented complaints about high frame rate videos (Ruppel et al., 2015; c.f. Wilcox et al., 2015). The relationship between visual motion smoothness and vection is an important question, given the relationship between vection and immersiveness (Prothero, 1998; Ohmi, 1998; Weech et al., 2019), but the nature of the association between frame rate and vection is currently understudied.

Recent work has established a monotonically-increasing relationship between frame rate and vection that asymptotes at 60 frames per second (FPS) (Fujii et al., 2019, 2018). A model was proposed by these authors linking visual motion energy and the sensation of vection, such that self-motion perception increases according to the amount of motion energy in a display, but asymptotes at high frame rates. These studies reported that the magnitude ratings, duration, and latency of vection tended to asymptote by 60 FPS, and often did so between 15-45 FPS. Fujii et al. interpreted values that approached this asymptote as ‘economical frame rates’, whereby a meaningful enhancement of vection would require a prohibitively large increase in FPS values. Other findings included evidence of increased vection for faster visual motion stimulation. On the other hand, the authors identified no significant difference in postural sway between 15-60 FPS displays, although there were major differences in the relationship between subjective (verbal) and objective (postural) vection measures for several participants. While these results indicate a possible asymptotic function that relates frame rate and vection, none of these experiments have employed frame rates greater than 60 FPS. Additionally, the two experiments reported by Fujii and co-workers (2018) provide evidence that the relationship between vection and frame rate depends on the characteristics of the visual stimulus: A downward-moving sine grating resulted in a higher asymptote for the function relating frame rate and vection when compared to a linearly expanding sine grating stimulus. This finding suggests the relevance of further examining this function for additional stimuli, such as those with well-defined luminance edges.

In the current study, we aimed to characterize the association between vection and frame rate across a wide range of FPS values between 15 and 480 FPS. To achieve highly-controlled high frame rate displays we used a DLP projection system with a scanning backlight, which has been developed to achieve deterministic timing for vision science applications. We assessed the magnitude, duration, and latency of vection, as well as excursions of the body centre of pressure (COP) as an objective measure of vection as a function of frame rates and the speed of visual motion for a linearly translating dot-cloud optic flow stimulus.

## Methods

### Participants

Twenty healthy participants (age *M* = 21.05 years, *SD* = 3.03 years, range = [18, 30], 15 female) were tested. All participants were pre-screened and reported no history of visual or vestibular disorders, musculoskeletal disorders, neurological conditions, or balance disorders. Written consent was obtained following written and verbal explanation of the study procedures and protocol. The study protocol was approved by the University of Waterloo ethics review board and adhered to the tenets of the Declaration of Helsinki.

### Design and Apparatus

Participants stood shoeless in tandem stance with their arms relaxed by their side on two force plates (4060-05; Bertec Corporation, Columbus, OH) with the right foot in front, one foot on each plate (Figure 1). The force plates were arranged one in front of the other with ~1 cm space in between. They were calibrated before data collection for each participant. The foot placement locations were marked on the plates with black tape. Participants stood facing a rigid rear-projection screen (acrylic with Da-Lite DA-100WA coating; 1.8 m × 1.0 m; 1.3 m throw distance from projector lens to screen) at 0.57 m viewing distance and were wearing lens-free goggles (80 × 50 deg visual field) that blocked the view of the edge of the screen. Participants were also given a computer mouse, held in their left hand, that was used to record when they experienced vection. The button on the mouse was to be pressed and held for the duration that the vection sensation was felt. Once no vection was felt the button was to be released.

**Fig 1.**
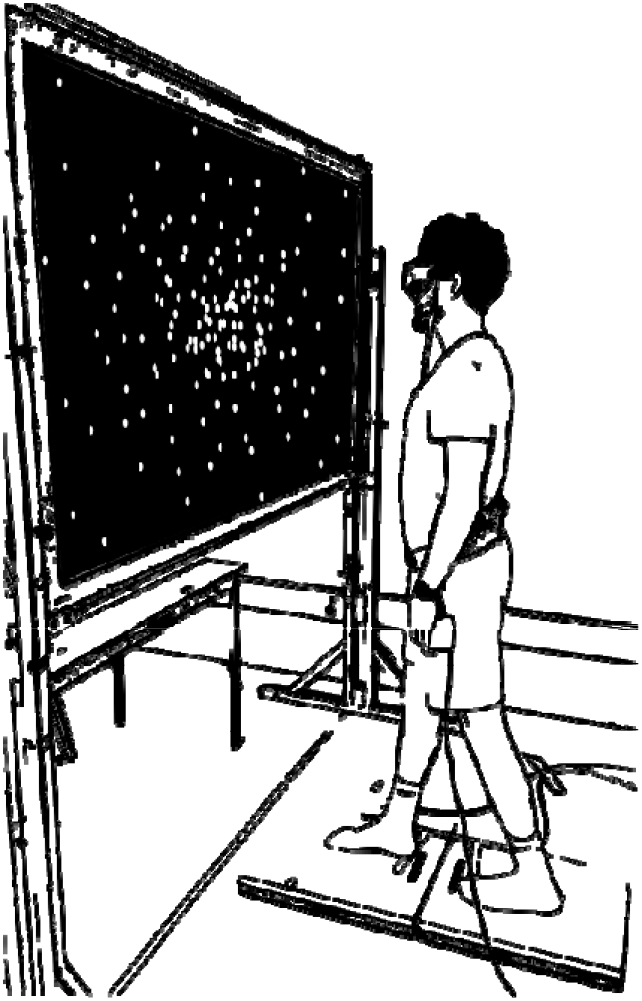
Depiction of experimental setup. Participants stood in tandem stance on two force plates in front of a projection screen while wearing lens free googles. Black tape on the force plates marked the location of right foot heel placement (front) and left foot toe placement (back). Goggles blocked the view of the edge of the screen. Participants held a computer mouse in their left hand to record the duration of felt vection.

Stimuli were rendered method in MATLAB (Mathworks Inc.) using Psychtoolbox (Brainard, 1997) and OpenGL rendering. The stimulus was back-projected onto the screen using a DLP LED projector (PROPixx, VPixx Technologies Inc., Montreal, Canada). The PROPixx operated in a specialized high refresh rate video sequencer mode called QUAD4X developed by VPixx Technologies (Alais et al., 2016; Balsdon et al., 2018; DeSimone & Schneider, 2019; Schweitzer & Rolfs, 2019; Zhang et al., 2019; Zhigalov et al., 2019). In short, the computer’s graphic card (Titan V, Nvidia, Santa Clara, CA) transmits a 1920 × 1080 pixels image at 120 FPS. Four frames are pre-rendered on individual quadrants of each of the full-resolution images. The PROPixx then presents these individual quadrants as four consecutive frames of 960 × 540 pixels, at 480 Hz. This results in updating the position of the optic flow stimuli every 2.1 ms.

All stimuli were displayed using the QUAD4X mode of the projector, and frame rates lower than 480 FPS were simulated by waiting a given number of frames within the stimulus generation program, before updating the position of the dots (e.g., 480 / 15 = 32; therefore, when simulating a 15 FPS refresh rate, the position of the dots is only updated every 32 frames while rendering at 480 FPS).

A custom-built LabVIEW program (National Instruments, Austin, TX) was used to record force plate data (vertical ground reaction forces and moments) for each 30 second trial. The online data was amplified using an internal digital preamplifier at a sampling rate of 1000 Hz. The data was then stored for off-line analysis. The force plate data was subsequently low-pass filtered (6-Hz, dual-pass 2nd-order Butterworth filter) and a second custom-made LabVIEW program was used to extract two postural sway features: The mean frequency of postural sway (Mean frequency), and the total path length of the COP throughout a trial (COP path length; Hufschmidt et al. 1980).

### Procedure

During each 30 second trial a radial optic flow stimulus–white dots on a black background–was displayed on the projection screen. Prior to each trial a fixation point (white) appeared on the screen for 3 sec, after which the cross disappeared. Participants were asked to maintain fixation on that location throughout the trial. There were 50 trials in total: 48 recorded and 2 ‘standard’ trials. The 48 recorded trials consisted of 6 levels of frame rate (spaced logarithmically: 15, 30, 60, 120, 240, and 480 FPS), and 2 levels of stimulus speed (slow [2 m/s] and fast [4 m/s]) which granted 12 unique trials; each of these 12 trials was repeated 4 times. Trials were carried out in a randomized order that was unique for each participant according to the method of constant stimuli. The ‘standard’ trial (30 FPS, slow) was presented to participants once at the beginning of the experiment (i.e., before the recorded trials) and once halfway after 24 recorded trials had elapsed.

Participants were asked to consider the amount of vection they felt during a standard trial as ‘100’. Vection was defined as the feeling of self-motion, which participants were told that they may or may not experience. Participants were told an example of vection was the sensation of self-motion felt when looking out of a stationary train window at a moving train. After each recorded trial the participant was asked to verbally rate their perceived vection relative to the vection that they experienced during the ‘standard’ stimulus. For example, after feeling twice the amount of vection as the standard the participant would respond ‘200’; for half, ‘50’; and for no vection, ‘0’. Participants could respond with any integer number they deemed representative of their relative sensation of vection. At the end of the experiment, participants were asked to report if they felt any motion sickness during or immediately after completing the task. This was intended to ensure consistently low levels of sickness across participants, given the known relationships between vection, presence, and motion sickness (Keshavarz et al., 2015; Weech et al., 2019). We conducted grand-mean centering for the unbounded vection measures (i.e., magnitude, mean frequency, and COP path length, according to Moskowitz, 1977).

## Results

All participants reported that the visual display evoked the sensation of vection during the experiment. No participants indicated any feelings of motion sickness due to exposure to the optic flow stimulus.

### Subjective Vection Measures

One-way analysis of variance (ANOVA) conducted on the factors of speed and FPS revealed significant main effects of both factors on the duration (FPS: *F*(5, 209) = 3.05, *p* = .011, η^2^ = 0.14, speed: *F*(5, 209) 52.77, *p* < .001, η^2^ = 0.73), magnitude (FPS: *F*(5, 209) = 3.46, *p* = .005, η^2^ = 0.15, speed: *F*(5, 209) = 204.21, *p* < .001, η^2^ = 0.92), and latency (FPS: *F*(5, 209) = 2.81, *p* = .017, η^2^ = 0.13, speed: *F*(5, 209) = 38.99, *p* < .001, η^2^ = 0.67) of vection. Data for the three subjective measures are plotted in Figure 2.

**Fig 2.**
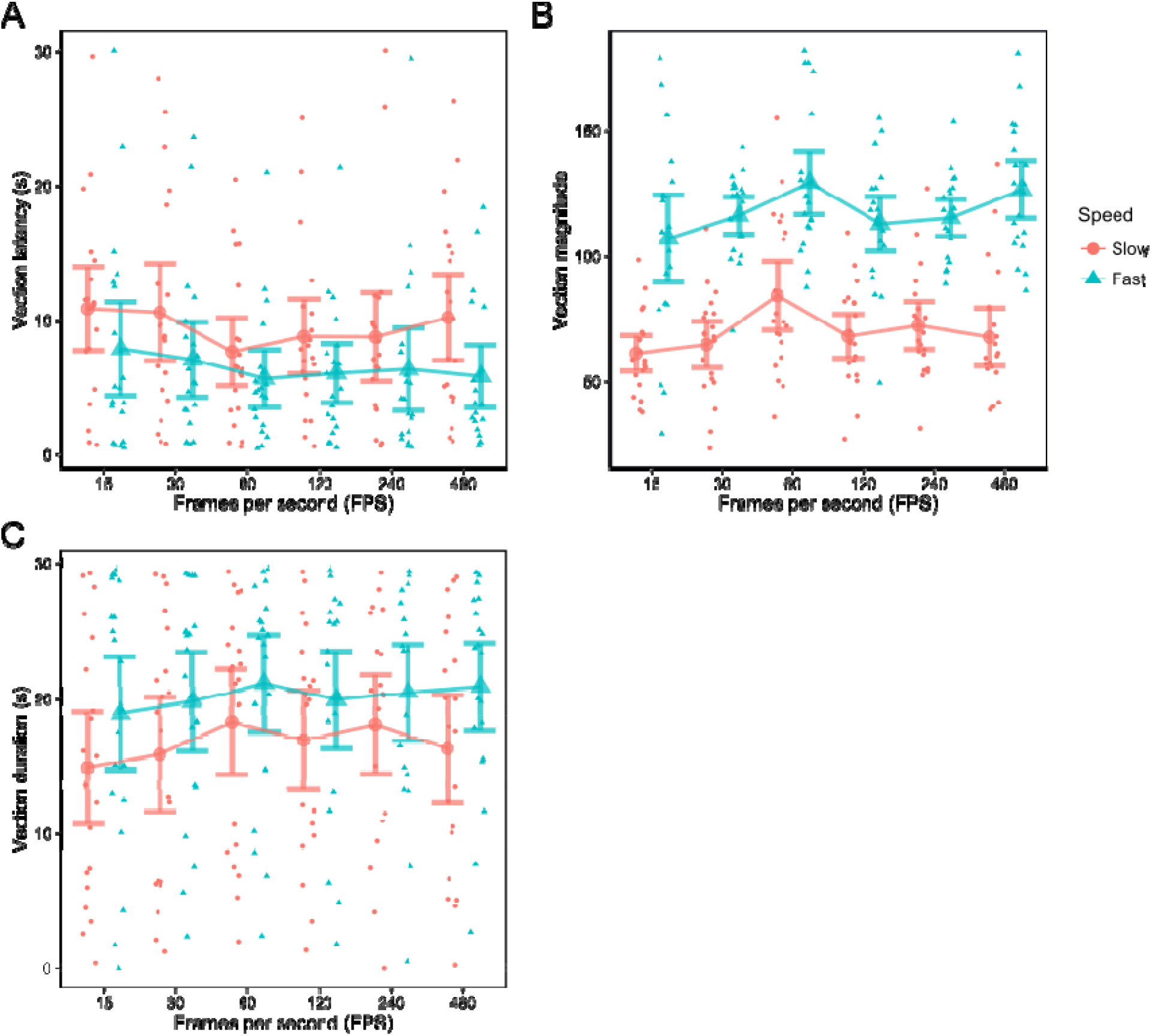
Line plots indicating average scores across FPS conditions for (A) latency, (B) magnitude, and (C) duration of vection. Each datapoint indicates a participant average. Red circles indicate slow speeds; Blue triangles indicate fast speeds. Error bars are 95% confidence intervals.

To further examine the main effects of speed we conducted *t*-tests examining the pairwise differences at each level of frame rate. For almost every condition we found that fast speed stimuli resulted in vection that was stronger magnitude, lower latency, and lasted longer (*p*s ≤ .044; except at 240 FPS for duration, where *p* = .054).

In order to characterise the main effects of FPS we conducted a second set of *t*-tests for pairwise differences at each level of speed. All significant comparisons for FPS conditions can be found in Table 1. First, we assessed if our results replicated the increase in vection from low frame rates (15 FPS) to high frame rates (60 FPS) found in previous research (Fujii et al., 2018, 2019). In line with these results, we saw a significant increase from 15 to 60 FPS in vection magnitude (slow speed: *t*(19) = 3.32, *p* = .004, Cohen’s *d* = 0.74; fast speed n.s.), duration (slow: *t*(19) = 3.13, *p* = .006, *d* = 0.70; fast: *t*(19) = 2.37, *p* = .029, *d* = 0.53), and decreased latency (slow: *t*(19) = −2.69, *p* = .014, *d* = 0.60; fast: *t*(19) = −2.19, *p* = .041, *d* = 0.49). We also found an increase from 30 FPS to 60 FPS in vection magnitude (*t*(19) = 2.80, *p* = .012, *d* = 0.63), duration (*t*(19) = 2.59, *p* = .018, *d* = 0.58), and a decrease in latency (*t*(19) = −3.41, *p* = .003, *d* = 0.76), all for slow speed stimuli.

**Table 1.** Significant (*p* < .05) pairwise comparisons for effects of FPS on subjective vection measures.

| Vection measure | Speed | Frame rates | $t(19)$ | $p$ | Cohen's $d$ |
| --- | --- | --- | --- | --- | --- |
| Magnitude | Slow | 60:15 | 3.32 | 0.004 | 0.74 |
|  | Slow | 240:15 | 3.04 | 0.007 | 0.68 |
|  | Slow | 60:30 | 2.80 | 0.012 | 0.63 |
|  | Fast | 480:15 | 2.74 | 0.013 | 0.61 |
|  | Slow | 480:60 | -2.36 | 0.029 | 0.53 |
|  | Slow | 120:60 | -2.15 | 0.045 | 0.48 |
|  | Fast | 480:120 | 2.12 | 0.048 | 0.47 |
| Duration | Slow | 60:15 | 3.13 | 0.006 | 0.70 |
|  | Slow | 60:30 | 2.59 | 0.018 | 0.58 |
|  | Slow | 240:15 | 2.38 | 0.028 | 0.53 |
|  | Slow | 480:60 | -2.37 | 0.028 | 0.53 |
|  | Fast | 60:15 | 2.37 | 0.029 | 0.53 |
|  | Fast | 120:60 | -2.31 | 0.032 | 0.52 |
| Latency | Slow | 60:30 | -3.41 | 0.003 | 0.76 |
|  | Slow | 60:15 | -2.69 | 0.014 | 0.60 |
|  | Slow | 480:60 | 2.50 | 0.022 | 0.56 |
|  | Fast | 60:15 | -2.19 | 0.041 | 0.49 |

*Note, positive $t$ -values indicate a greater value for first FPS condition and vice versa.*

Our most novel results related to frame rates above 60 FPS that have not been examined in previous studies. We observed an increase from 15 to 240 FPS in vection magnitude (*t*(19) = 3.04, *p* = .007, *d* = 0.68) and duration (*t*(19) = 2.38, *p* = .028, *d* = 0.53) for slow stimuli. Additionally, when compared with 60 FPS, we found that vection for 480 FPS stimuli was lower in magnitude (*t*(19) = −2.36, *p* = .029, *d* = 0.53), duration (*t*(19) = −2.37, *p* = .028, *d* = 0.53), and had increased latency (*t*(19) = 2.50, *p* = .022, *d* = 0.56), all for slow speed stimuli. In fact, 60 FPS reliably resulted in greater vection than most other conditions, including 120 FPS (magnitude: *t*(19) = −2.15, *p* = .045, *d* = 0.48, slow; duration: *t*(19) = −2.31, *p* = .032, *d* = 0.52, fast). Finally, for fast stimuli, vection magnitude was greater for 480 FPS compared to 120 FPS stimuli (*t*(19) = 2.12, *p* = .048, *d* = 0.47) and 15 FPS stimuli (*t*(19) = 2.74, *p* = .013, *d* = 0.61).

### Objective Vection Measures

For the objective correlates of vection we ran ANOVAs on the factors of FPS and speed for mean frequency and COP path length (see Figure 3 for data). A significant main effect of speed emerged for Mean Frequency (*F*(1,235) = 7.06, *p* = .008, η^2^ = 0.16), and follow-up *t*-tests revealed that this effect was driven by a higher frequency at low FPS (15) (*t*(19) = −2.88, *p* = .0097, *d* = 0.87) for slow speeds. There was no effect of FPS on mean frequency (*F*(1,235) = 0.04, *p* = .845, η^2^ = 0.01) and no interaction between FPS and speed (*F*(1,235) = 0.15, *p* = .698, η^2^ = 0.04).

**Fig 3.**
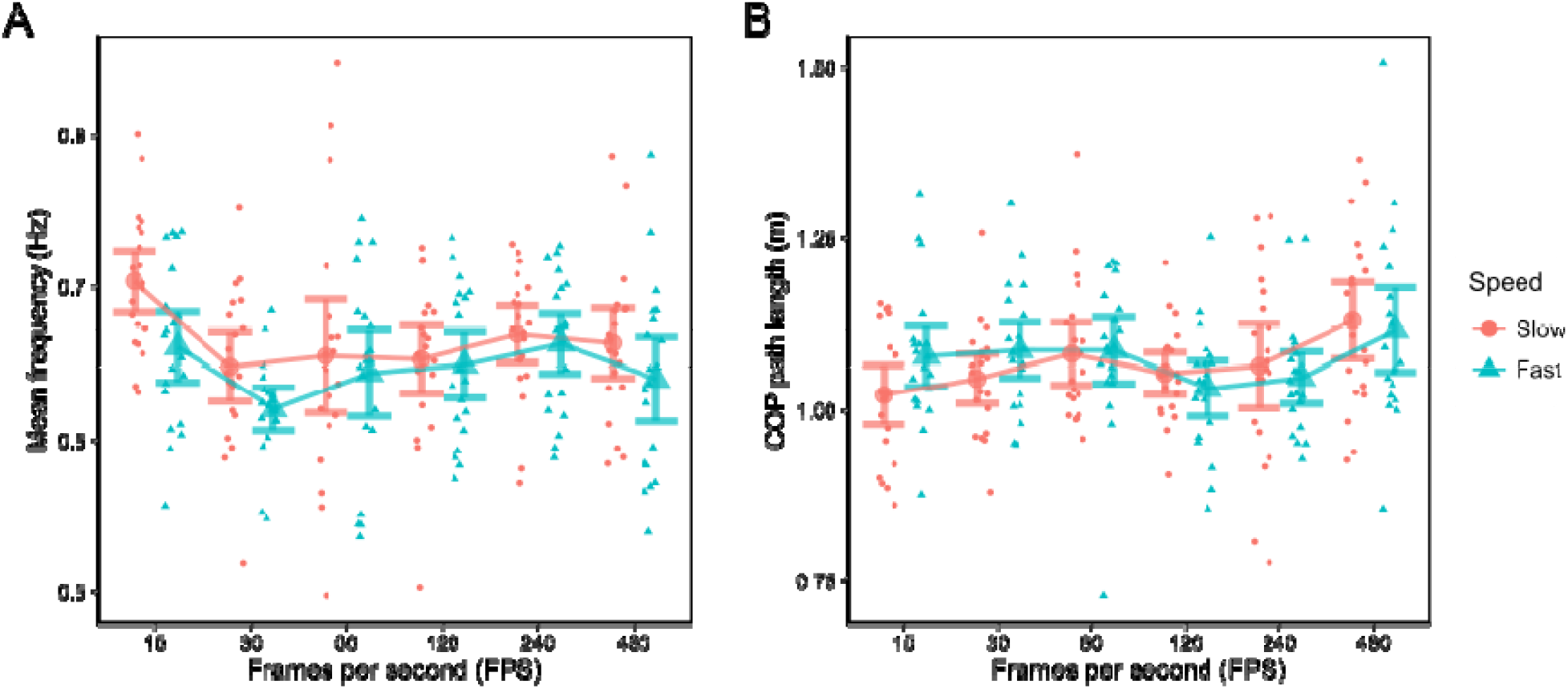
Line plots indicating average scores across FPS conditions for (A) mean frequency of sway and (B) COP path length. Each datapoint indicates a participant average. Red circles indicate slow speeds; Blue triangles indicate fast speeds. Error bars are 95% confidence intervals.

We also obtained a significant main effect of FPS for COP path length (*F*(1,235) = 8.00, *p* = .005, η2 = 0.17). Follow-up *t*-tests revealed that this effect was driven by a higher COP for 480 FPS stimuli when compared to 15 FPS (*t*(19) = 3.32, *p* = .004, *d* = 0.74), 30 FPS (*t*(19) = 2.20, *p* = .041, *d* = 0.49), and 120 FPS (*t*(19) = 2.20, *p* = .041, *d* = 0.49) at slow speeds. We found no effect of speed (*F*(1,235) = 0.38, *p* = .539, η^2^ = 0.01) and no speed-FPS interaction (*F*(1,235) = 1.98, *p* = .161, η^2^ = 0.05).

### Subjective and Objective Vection Correlation

Correlations between objective and subjective vection were found only at low frame rates. We observed a negative correlation between vection duration and mean frequency for 15 FPS at low speeds (*r*(18) = −.502, *p* = .024), and a positive correlation between latency and mean frequency for the same condition (*r*(18) = .624, *p* = .003). For fast speed stimuli, we found significant positive correlations between vection magnitude and both mean frequency (15 FPS; *r*(18) = .504, *p* = .023) and COP path length (*r*(18) = .467, *p* = .037). All other correlations between objective and subjective indices were non-significant (*p*s > .05).

### Exponential asymptotic curve fitting

We fit the data to exponential curves according to the function proposed by Fujii and colleagues (2018) (*y = a*exp(−b*x)+c* for the fit parameters a, b, and c) and found a large divergence between our results and the previous data on several counts. While some of our data are well represented by this equation (e.g., latency data for fast stimuli), overall the data do not conform to an exponential function. For model fit evaluation, we report the *R*^2^ goodness of fit for each fitted equation as well as the adjusted *R*^2^ that depicts fit while accounting for model complexity (values not reported in Fujii et al. 2018). As a comparison, we then use these curves to compute ‘economical frame rates’ (i.e., within a 95% CI of the estimated asymptote; Fujii et al. 2018), although since our data are often not well fitted by these exponential curves, these computed values should be treated conservatively. Economical frame rates are intended to depict the level above which increasing FPS would require significant computational power (e.g., graphical resources) to achieve a meaningful increase in vection. *R*^2^, adjusted *R*^2^, economical frame rates, and function parameters are depicted in Table 2.

**Table 2.** Parameters and goodness-of-fit measures for exponential functions (*y = a*exp(−b*x)+c*).

| Vection measure | Speed | $R^2$ ($R^2_{Adj}$) | a | b | c | EFR |
| --- | --- | --- | --- | --- | --- | --- |
| Magnitude | Slow | 35.85 (-7.92) | -37.03 | 0.075 | 72.12 | 31.06 |
|  | Fast | 43.25 (5.4) | -60.88 | 0.098 | 120.40 | 23.59 |
| Latency | Slow | 45.36 (8.93) | 5.09 | 0.058 | 8.89 | 41.92 |
|  | Fast | 87.28 (78.8) | 4.64 | 0.057 | 5.92 | 47.94 |
| Duration | Slow | 62.49 (37.48) | -6.81 | -6.810 | 17.35 | 32.44 |
|  | Fast | 69.78 (49.63) | -5.02 | -5.020 | 20.58 | 22.39 |
| Mean frequency | Slow | 85.54 (75.9) | 13.98 | 13.980 | 0.66 | 17.80 |
|  | Fast | 20.99 (-31.68) | 73.00 | 0.556 | 0.64 | 14.25 |
| COP path length | Slow | 34.02 (-9.96) | -23.12 | 0.406 | 1.07 | 19.56 |
|  | Fast | - | - | - | - | - |

*note, EFR = Economical frame rate (Fujii et al., 2018)*

### Economical frame rates: Subjective measures

Economical frame rates for our stimuli were on average 35.14 FPS (slow speed) and 31.31 FPS (fast speed) pooling across subjective vection measures; and 27.33 for vection magnitude, 44.93 for latency, and 27.42 for duration when pooling across speeds.

### Economical frame rates: Objective measures

As shown in Table 1, objective vection parameters were generally poorly fit by an exponential function. There was no solution for a fit to the exponential curve for COP path length for fast speed stimuli. On the other hand, we found a relatively good exponential fit between FPS and mean frequency for slow speeds, which demonstrated the highest *R*^2^ and adjusted *R*^2^ values. While the average economical frame rates suggested by our fitting procedure for mean frequency (16.03 FPS) and COP path length (19.56 FPS) were very low, the lack of a robust fit to the proposed exponential function precludes a strong reliance on this result.

## Discussion

Here we set out to examine the relationship between vection and frame rate of a visual display. Based on previous evidence of an asymptotic exponential relationship at frame rates up to 60 FPS, we expected to find no differences between vection measures at 120, 240, or 480 FPS. This was not the case; While 60 FPS was superior in evoking subjective vection compared to higher frame rates, a 480 FPS display was more effective in generating objective vection as measured by COP path length excursions.

Considering the high computational demands of high frame rate video with respect to information bandwidth and graphical rendering power, the superiority of 60 FPS video in evoking stronger, longer lasting, and earlier vection recommends the use of this frame rate in applications intended to evoke a compelling sense of self-motion. At the same time, we observed the greatest level of vection (objective: COP path length) for the 480 FPS stimuli at slow speeds, which was significantly higher than 15, 30, and 120 FPS stimuli, which supports the utility of ultra-high frame rate stimuli for inducing vection. How valuable is this increase? We would contend that it is relatively low, given the significant computing power required to achieve ultra-high frame rate. Despite this, rapid progression in graphics technology (e.g., deep-learning supported 3D graphics processing; Bui et al., 2018; Nalbach et al., 2017) means the capability for high-resolution, high frame rate rendering will soon be widely available, such that even small enhancements in vection will be easily attainable. However, the lack of improvement in *subjective* vection for high FPS displays suggests that values of around 15-45 FPS, around the economical frame rates of our study and others (Fujii et al., 2018), are optimal for inducing compelling self-motion, at least for the impoverished stimuli evaluated in these experiments.

Across the measures of vection we obtained, our data do not adhere to a simple exponential function. This result contrasts with those of previous research, who identified asymptotic limits for vection at approximately 15-45 FPS. Rather, we have shown two pieces of evidence that contradict the idea of an asymptote around this level. First, we found that 60 FPS results in superior subjective vection than all other frame rates tested for slow speed stimuli. Second, we saw an increase in objective vection that peaked at 480 FPS for slow speed stimuli.

The results obtained here show that 60 FPS stimuli provoked stronger vection than all other frame rates. As yet, there is no apparent reason for a preference for 60 FPS in generating a compelling sense of self-motion over and above higher frame rates. If motion energy is the major visual factor underlying vection (Fujii et al., 2018), the highest frame rate stimuli should result in the strongest vection; while we found this result for objective vection, it was not so for subjective vection. The most “familiar” visual displays would likely have been those closest to natural stimuli (which have “infinite” refresh rate). For this reason, previous evidence that familiar types of stimuli (e.g., more realistic scenes, more familiar places) evoke stronger vection (Bubka & Bonato, 2010; Nakamura, 2010; Riecke et al., 2006; Van der Steen & Brockhoff, 2000) would suggest that 480 FPS should produce the strongest vection. Since no previous study has implemented frame rates of up to 480 FPS in the context of vection research, this result is novel, and it provokes further investigation.

While our data are not well fit by an exponential function, the results with respect to economical frame rates are similar to those of previous research (Fujii et al., 2018). This is the case despite a strong dependency of these values on visual motion characteristics, as evidenced by differences of around 20-30 FPS in economical frame rates across Experiments 1 and 2 for the previous study (a downwards-moving sine grating, and a radially expanding sine-grating, respectively). Given our estimated economical frame rates, we agree with their suggestion that traditional movie content (usually around 24 FPS) are likely to be insufficient for evoking the economically optimal level of vection (up to 48 FPS in our study). At the same time, other perceptual aspects must be considered in the context of evaluating ‘optimality’ for video frame rates. For instance, familiarity appears to be a strong motivator of the preference for low frame rate movies compared to high frame rate movies (Wilcox et al., 2015). Moreover, the cost-benefit analysis for cinema projection systems is vastly different to the analysis for real-time graphical rendering (e.g., in virtual reality), where computing resources are often highly limited. The utility of high frame rate video must therefore be evaluated according to context, and the strength of evoked vection is but one aspect of this decision.

We found only moderate evidence of correlations between subjective and objective measures of vection. We observed correlations between subjective and objective vection at low frame rates (15/30 FPS). COP path length and vection magnitude were only correlated at 30 FPS for fast speeds, where high total COP sway was associated with high vection magnitude. At 15 FPS, we found a mixture of results: high frequency sway was associated with high vection magnitude, but also high vection latency and low vection duration. All other correlations between objective and subjective indices were non-significant. While the lack of strong correlation between the two measurement classes was unexpected, discrepancies between postural and subjective measures of vection have been reported before (Delorme & Martin, 1986; Fujii et al., 2019; Kawakita et al. 2000; Palmisano et al., 2014; Weech et al., 2018). Our finding reiterates the need to concurrently evaluate objective and subjective vection indices in studies such as these, as evidence is accruing that there may be a major dissociation in the results obtained by these two approaches to measuring the ‘same’ percept.

It should be noted that the visual stimuli used in our study were relatively simplistic, consisting of a dot cloud floating in an abyss. This simplicity was intentional in order to maintain stable frame timing at high frame rates. A more naturalistic visual stimulus might result in a different pattern of results given that scene realism modulates vection, with more realistic scenes generating a stronger sensation of self-motion (Bubka & Bonato, 2010; Nakamura, 2010; Riecke et al., 2006; Van der Steen & Brockhoff, 2000). While it is not yet possible to generalize our results to realistic visual stimuli, previous research has confirmed the existence of a monotonically increasing relationship between frame rate and vection for high-fidelity virtual scenes (Fujii et al., 2019). Additional replication studies will be able to take advantage of future developments in graphical technology to present highly realistic, ultra-high frame rate stimuli with stable frame timing.

## Conclusions

Our results support the existence of diminishing returns for vection at high frame rates. At the same time, the relationship between frame rate and vection is not well represented by an asymptotic exponential function according to our data; rather, dependencies emerge between stimulus speed, vection measurement approach, and frame rate when testing vection at ultra-high frame rates.

